# SARS-CoV-2 envelope protein induces LC3 lipidation via the V-ATPase-ATG16L1 axis

**DOI:** 10.1101/2025.06.12.659257

**Authors:** Carmen Figueras-Novoa, Lewis Timimi, Elena Marcassa, Raquel Taveira-Marques, Lorin Adams, Ming Jiang, Mary Wu, Beatriz Montaner, Kevin Ng, Giuditta De Lorenzo, Wilhelm Furnon, Vanessa M. Cowton, Nicole Upfold, George Kassiotis, Ruth Harvey, Arvind H Patel, Michael Howell, Rachel Ulferts, Rupert Beale

## Abstract

Coronaviruses encode envelope (E), a structural component of the virion that is important for assembly and egress. E has proton channel activity that prevents premature rearrangement of the spike glycoprotein as virions encounter acidic compartments as they exit the cell. How infected cells respond to this pH disruption during coronavirus infection is unknown. Here we show that SARS-CoV-2 E ion channel activity triggers the proton pump V-ATPases to recruit the ATG16L1 complex during infection. This results in ATG8 molecules such as LC3B decorating perturbed compartments. This recruitment of autophagy machinery does not inhibit viral replication, rather SARS-CoV-2 exploits this response. Inhibition of the V-ATPase/ATG16L1 interaction using the *Salmonella* effector SopF inhibits SARS-CoV-2 replication. Careful distinction between use of the autophagic machinery from canonical macroautophagy is required in order to better understand coronavirus replication and for rational targeting of any potential host-directed therapies.

## Introduction

Viruses generally enter cells by binding to a surface receptor and becoming internalised by endocytosis (Yamauchi and Helenius, 2013). Acidification of endosomes frequently acts as a signal for the virus to disrupt the endosomal membrane and invade the cytosol (Mercer et al., 2010, White and Whittaker, 2016). This creates an important biological problem which a successful virus must solve: to create infectious virions, the viral entry proteins must first avoid acidic conditions where premature triggering of conformations required for fusion might occur. The secretory pathway is acidic (Linders et al., 2022), so viruses that employ this pathway to assemble structural components must subvert host cell physiology to avoid creating inert virions incapable of invading a new host cell.

Coronaviruses employ their spike glycoprotein (S) to bind to a suitable receptor for cell entry (Jackson et al., 2022). S then undergoes a conformational change at low pH to expose hydrophobic regions that leads to fusion of the viral and endosomal membranes, allowing entry of the viral genome to the cytosol where replication takes place (Jackson et al., 2022). Coronaviruses employ envelope (E) a virally encoded ion channel that prevents acidification of secretory compartments by draining protons from the Golgi and post-Golgi compartments to protect S (Westerbeck and Machamer, 2019, Cabrera-Garcia et al., 2021). We have shown that the Influenza A virus (IAV) ion channel Matrix 2 (M2), that plays a parallel role in protecting IAV haemagglutinin by deacidifying the secretory pathway (Ciampor et al., 1992), induces a cellular response to de-acidification that superficially resembles autophagy (Beale et al., 2014). When cellular compartments cannot be correctly acidified, the proton pump vacuolar ATPase (V-ATPase) itself directly recruits components of the autophagy machinery (Ulferts et al., 2021, Timimi et al., 2024). Inactive, fully assembled V-ATPases accumulate following pH gradient disruption and recruit the E3 ubiquitin-ligase like ATG5–ATG12-ATG16L1 complex that covalently lipidates ubiquitin-like ATG8 molecules to tag the membrane of deacidified compartments (Timimi et al., 2024). Lipidation of ATG8 molecules to double membrane autophagosomes is commonly used as a marker for macroautophagy, a process by which cytosolic contents are targeted for lysosomal degradation. However, V-ATPase mediated recruitment of ATG5–ATG12-ATG16L1 leads to conjugation of ATG8s at single membranes (CASM), independently of upstream autophagy regulators (Florey et al., 2011, Ulferts et al., 2021, Figueras-Novoa et al., 2024).

V-ATPase-ATG16L1 Induced LC3 lipidation (VAIL) has been shown to play an important role in the response to intracellular infection. Stimulation of the innate immune sensor STING allows it to acquire proton channel activity, leading to VAIL (Liu et al., 2023, Fischer et al., 2020). Furthermore, numerous bacterial pathogens induce VAIL through neutralisation of organelle pH, including *Salmonella* Typhimurium (Xu et al., 2019), *Listeria monocytogenes* (Gluschko et al., 2018), *Mycobacterium tuberculosis* (Köster et al., 2017), and *Shigella* flexneri (Xu et al., 2019, Campbell-Valois et al., 2015). IAV triggers VAIL through the activity of the virally encoded proton channel M2 (Fletcher et al., 2018, Ulferts et al., 2021, Ren et al., 2016). Disruption of M2-induced VAIL alters the morphology and reduces the stability of IAV virions (Beale et al., 2014, Figueras-Novoa et al., 2025), while VAIL-deficient mice exhibit increased lung inflammation and mortality when infected with IAV (Wang et al., 2021).

SARS-CoV-2 is an enveloped, single stranded RNA virus from the genus *Betacoronavirus* of the family *Coronaviridae* (Zhou et al., 2020). The SARS-CoV-2 pandemic has led to millions of deaths (Veiga and Cavalcanti, 2023). Given the clinical significance of this pathogen, there is a clear need to understand the mechanisms by which SARS-CoV-2 interacts with and subverts host pathways during replication and in disease.

The induction of LC3 lipidation during infection with SARS-CoV-2 is well established (Gordon et al., 2020, Gassen et al., 2021, Miao et al., 2021, Shang et al., 2021, Qu et al., 2021, Gorshkov et al., 2021). However, no studies have explored the contribution of VAIL to these processes. Through a combination of genetic tools designed to distinguish autophagy and VAIL, we reveal the ability of E to induce VAIL during SARS-CoV-2 infection and its importance for viral replication.

## Results

### SARS-CoV-2 infection induces VAIL

Multiple reports have described the induction of LC3 lipidation during SARS-CoV-2 infection, but to date none have examined the contribution of VAIL to the cellular pool of lipidated LC3 (Gordon et al., 2020, Gassen et al., 2021, Miao et al., 2021, Qu et al., 2021, Shang et al., 2021, Gorshkov et al., 2021). Infection of A549 ACE2/TMPRSS2 cells with SARS-CoV-2 (England/02/2020) resulted in robust and reproducible LC3B lipidation (Figures 1A and 1B). Furthermore, imaging of A549 ACE2/TMPRSS2 cells expressing GFP-LC3B demonstrated GFP-LC3B puncta formation during infection, confirming the induction of LC3 lipidation (Figure 1C). As VAIL requires the V-ATPase complex for initiation (Florey et al., 2015, Hooper et al., 2022, Timimi et al., 2024), we also analysed the distribution of the V-ATPase subunit V1D by immunofluorescence microscopy. This revealed that multiple GFP-LC3B-positive structures within infected cells were also positive for the V1D subunit (Figure 1D), suggesting that the V-ATPase may be involved in the initiation of LC3 lipidation during SARS-CoV-2 infection.

**Figure 1:**
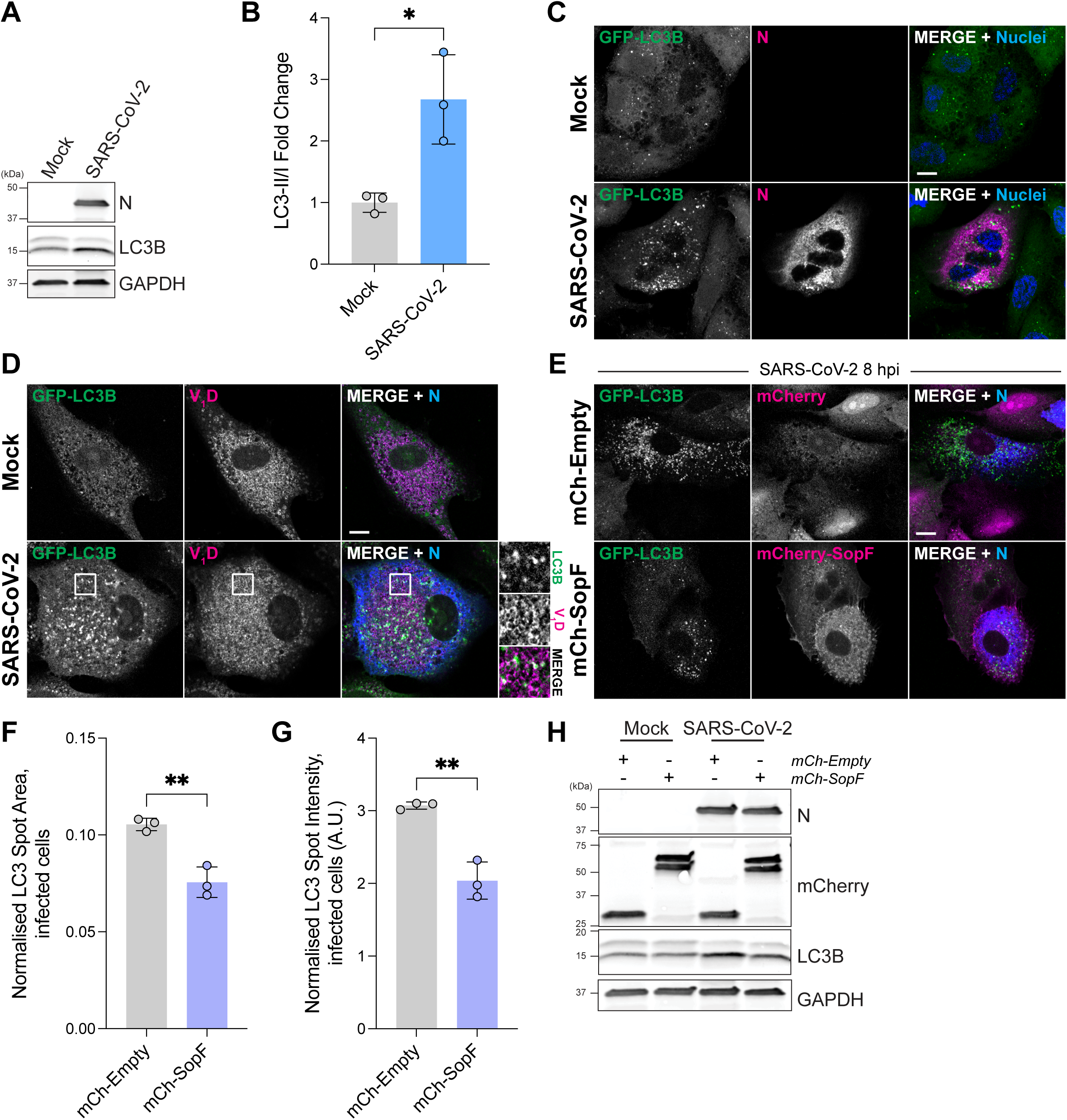
SARS-CoV-2 infection induces VAIL. A. Western blot of A549 cells expressing ACE2/TMPRSS2. Indicated cells were infected with SARS-CoV-2 (MOI 1) and lysed at 16 h post infection (hpi). B. Quantification of A. Bars show mean ± SD; n=3. *: P ≤ 0.05. Unpaired *t*-test. C. Representative images of A549 cells expressing ACE2/TMPRSS2 and EGFP-LC3B (in green). Indicated cells were infected with SARS-CoV-2 (MOI 5), fixed at 16 h post infection (hpi) and stained with SARS-CoV-2 nucleocapsid (N; in magenta) and NucBlue (nuclei; in blue). Scale bar, 10 µm. D. Representative images of A549 cells expressing ACE2/TMPRSS2 and EGFP-LC3B (in green). Indicated cells were infected with SARS-CoV-2 (MOI 5), fixed at 16 h post infection (hpi) and stained with V1D (V1D; in magenta) and SARS-CoV-2 nucleocapsid (N; in blue). Scale bar, 10 µm. E. Representative images of A549 cells expressing ACE2/TMPRSS2, EGFP-LC3B (in green), and either mCherry-Empty or mCherry-SopF (as indicated; in magenta). Cells were infected with SARS-CoV-2 (MOI 5), fixed at 16 h post infection (hpi) and stained with SARS-CoV-2 nucleocapsid (N; in blue). Scale bar, 10 µm. F. Quantification of EGFP-LC3B relocalisation in A549 cells expressing ACE2/TMPRSS2, EGFP-LC3B and either mCherry-Empty or mCherry-SopF. Cells were infected with SARS-CoV-2 (MOI 2), fixed at 24 h post infection (hpi), stained with SARS-CoV-2 nucleocapsid and analysed using automated high-content imaging. The total area of EGFP-LC3B spots within infected cells was normalised to the total nucleocapsid-positive area in each well. Bars show mean ± SD; n=3. **: P ≤ 0.01. Unpaired *t*-test. G. Further quantification from F. The total intensity of EGFP-LC3B spots within infected cells was normalised to the total nucleocapsid-positive area in each well. Bars show mean ± SD; n=3. **: P ≤ 0.01. Unpaired *t*-test. H. Western blot of A549 cells expressing ACE2/TMPRSS2 and either mCherry-Empty or mCherry-SopF. Indicated cells were infected with SARS-CoV-2 (MOI 5) and lysed at 16 h post infection (hpi).

To investigate the role of VAIL directly, we utilised the *Salmonella* effector protein SopF. SopF ADP-ribosylates the V-ATPase complex to block V-ATPase-ATG16L1 binding, resulting in the inhibition of VAIL (Xu et al., 2019). Importantly, SopF has no effect on LC3 lipidation after multiple different stimuli that induce canonical macroautophagy (Xu et al., 2019, Ulferts et al., 2021). A549 ACE2/TMPRSS2 cells expressing GFP-LC3B were transduced with either mCherry-Empty vector or mCherry-SopF. Imaging of these cells after infection with SARS-CoV-2 showed a reduction in GFP-LC3B relocalisation in mCherry-SopF cells relative to mCherry-Empty cells (Figure 1E). Automated image acquisition and analysis confirmed a reduction in LC3B punctum area and intensity within infected cells expressing mCherry-SopF compared to mCherry-Empty (Figures 1F and 1G). Western blot detection of LC3 verified the reduction of infection-induced LC3 lipidation in SopF-expressing cells (Figure 1H). Together, these results indicate that LC3 lipidation induced by SARS-CoV-2 infection is in part mediated by VAIL.

### Inhibition of VAIL attenuates SARS-CoV-2 replication

While assessing the impact of SopF expression on SARS-CoV-2 infection, we noticed that SopF-expressing cells contained a reduced amount of nucleoprotein (N) protein at late timepoints (Figure 2A). To explore this observation, we performed RT-qPCR of supernatants from infected mCherry-Empty and mCherry-SopF cells at 24 h post infection (h p.i.). Copy numbers of both SARS-CoV-2 N and RNA-dependent RNA polymerase (RdRp) in supernatant from SopF-expressing cells were reduced (Figures 2B and 2C). Automated imaging revealed a decrease in the number of N-positive cells in SopF-expressing cells infected with SARS-CoV-2 for 24 hours when compared to mCherry-empty cells (Figures 2D and 2E). This decrease in infection was observed despite comparable mCherry levels across mCherry-empty and mCherry-SopF cells (Figure S1A). Furthermore, N-positive cells across SopF and control cells expressed equivalent N intensity, suggesting a defect in viral exit and no effect viral protein production (Figure S1B). Assessment of virus production by plaque assay also revealed a log-fold reduction in titre from the supernatant of virus cultured for 24 h in SopF-expressing A549 ACE2/TMPRSS2 cells (Figure S1C). Taken together, these results point to a disruption in SARS-CoV-2 replication within VAIL-deficient cells.

**Figure 2:**
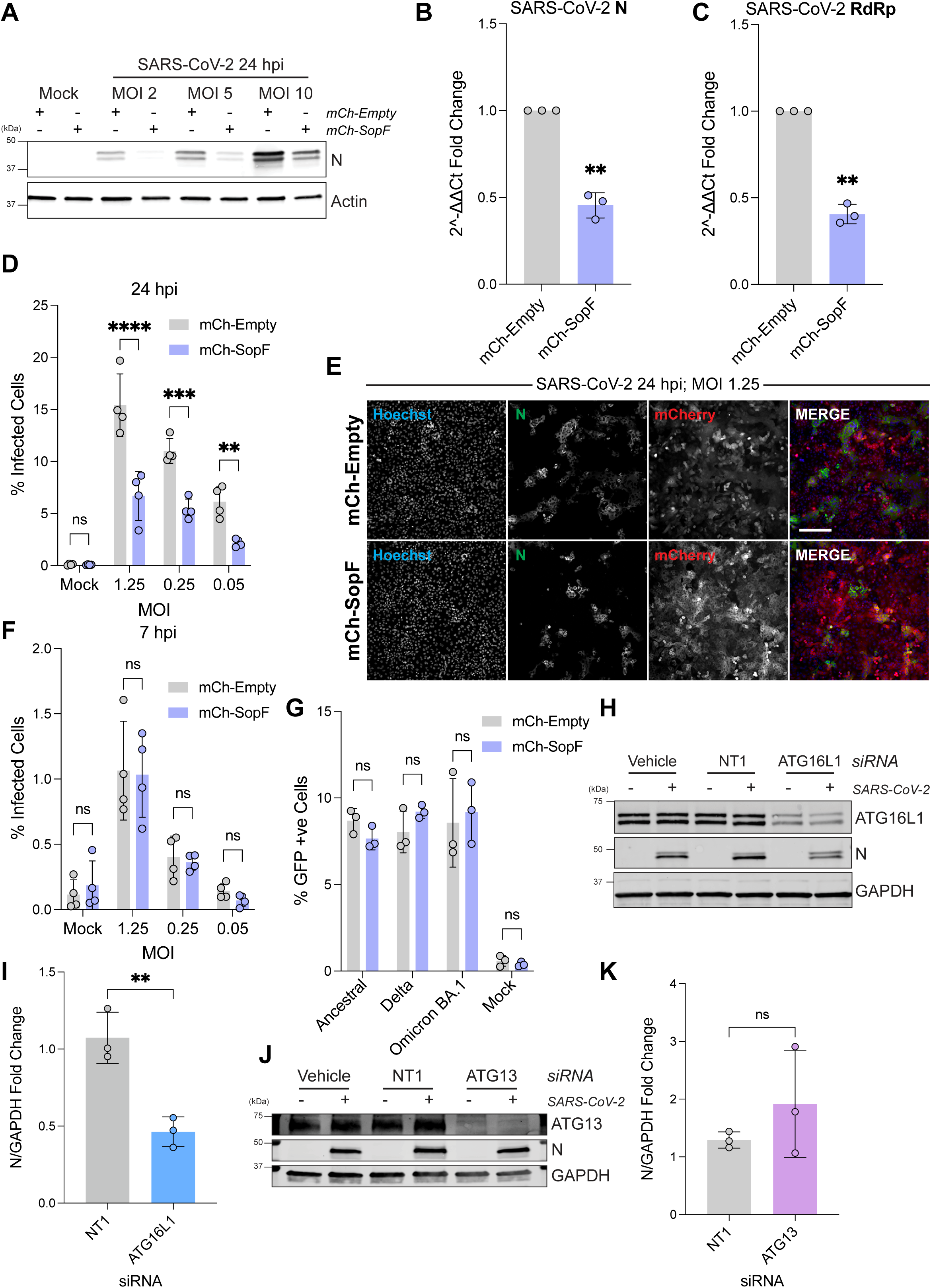
Inhibition of VAIL attenuates SARS-CoV-2 replication A. Western blot of A549 cells expressing ACE2/TMPRSS2. Cells were infected with SARS-CoV-2 at the indicated MOI and lysed 24 h post infection (hpi). B. RT-qPCR analysis of SARS-CoV-2 nucleocapsid (N) from A549 cells expressing ACE2/TMPRSS2 and either mCherry-Empty or mCherry-SopF. Cells were infected with SARS-CoV-2 (MOI 2) and lysed at 24 h post infection (hpi). Bars show mean ± SD; n=3. **: P ≤ 0.01. One-sample *t*-test. C. RT-qPCR analysis of SARS-CoV-2 RNA-dependent RNA polymerase (RdRp) from A549 cells expressing ACE2/TMPRSS2 and either mCherry-Empty or mCherry-SopF. Cells were infected with SARS-CoV-2 (MOI 2) and lysed at 24 h post infection (hpi). Bars show mean ± SD; n=3. **: P ≤ 0.01. One-sample *t*-test. D. Percentage of infected (nucleocapsid-positive) A549 cells expressing ACE2/TMPRSS2 and either mCherry-Empty or mCherry-SopF. Cells were infected with SARS-CoV-2 at the indicated MOI (MOIs were based on titres determined by Vero E6 plaque assay). Plates were fixed at 24 h post infection (hpi), stained with SARS-CoV-2 nucleocapsid and analysed using automated high-content imaging. Bars show mean ± SD; n=4. **: P ≤ 0.01; ***: P ≤ 0.001; ****: P ≤ 0.0001. Two-way ANOVA with Šídák’s multiple comparisons. E. Representative image from D. Scale bar, 200 µm. F. Percentage of infected (nucleocapsid-positive) A549 cells expressing ACE2/TMPRSS2 and either mCherry-Empty or mCherry-SopF. Cells were infected with SARS-CoV-2 at the indicated MOI, fixed at 7 h post infection (hpi), stained with SARS-CoV-2 nucleocapsid and analysed using automated high-content imaging. Bars show mean ± SD; n=4. Two-way ANOVA with Šídák’s multiple comparisons. G. Transduction of A549 cells expressing ACE2/TMPRSS2 and either mCherry-Empty or mCherry-SopF using GFP lentiviral particles pseudotyped with SARS-CoV-2 Spike from the indicated lineages. The percentage of transduced (GFP-positive) cells was assessed by flow cytometry at 72 h post transduction. Bars show mean ± SD; n=3. Two-way ANOVA with Šídák’s multiple comparisons. H. Western blot of A549 cells expressing ACE2/TMPRSS2 following knockdown of ATG16L1. NT1 refers to non-targeting control siRNA. Indicated cells were infected with SARS-CoV-2 (MOI 2) and lysed at 16 h post infection (hpi). I. Quantification of N/GAPDH ratio from infected samples in H, relative to vehicle only. Bars show mean ± SD; n=3. **: P ≤ 0.01. Unpaired *t*-test. J. Western blot of A549 cells expressing ACE2/TMPRSS2 following knockdown of ATG13. NT1 refers to non-targeting control siRNA. Indicated cells were infected with SARS-CoV-2 (MOI 2) and lysed at 16 h post infection (hpi). K. Quantification of N/GAPDH ratio from infected samples in J, relative to vehicle only. Bars show mean ± SD; n=3. Unpaired *t*-test.

Interestingly, the difference in infection levels observed with SopF was not seen at earlier timepoints, suggesting that virus entry was unaffected by the inhibition of VAIL (Figure 2F). To examine the impact of SopF on viral entry, we performed pseudotyped virus infection assays using GFP lentiviral particles bearing SARS-CoV-2 spike receptors. Infection of cells with all SARS-CoV-2 pseudoviruses tested was unaffected by the presence of SopF (Figure 2G), indicating that the impact of VAIL inhibition on SARS-CoV-2 replication is likely due to defects in later stages of the viral life cycle.

To verify that the effects of SopF on SARS-CoV-2 replication were related to the inhibition of VAIL, we tested the requirement for other VAIL components in SARS-CoV-2 replication. VAIL requires the autophagic ATG8 lipidation machinery, but not the upstream autophagic initiation complexes (Xu et al., 2019, Ulferts et al., 2021, Hooper et al., 2022); therefore, inhibition of VAIL occurs upon disruption of the lipidation machinery, but not upstream autophagy regulators. In line with this, knockdown of ATG16L1 in ACE2/TMPRSS2 cells led to a reduction in the abundance of N in SARS-CoV-2 infected cells (Figures 2H and 2I). Crucially, knockdown of ATG13, a component of the ULK complex required for autophagic initiation (Jung et al., 2009), produced little effect on infection (Figures 2J and 2K). Thus, disruption of VAIL attenuates SARS-CoV-2 replication.

### SARS-CoV-2 envelope (E) induces VAIL

Having identified a role for VAIL in SARS-CoV-2 replication, we sought to understand how the virus induces this cellular pathway. VAIL is typically triggered by the neutralisation of physiologically acidified organelles within cells (Durgan and Florey, 2022, Figueras-Novoa et al., 2024). The E proteins of coronaviruses, including SARS-CoV-2, form cation-conducting ion channels that neutralise Golgi and endolysosomal pH during infection (Mandala et al., 2020, Cabrera-Garcia et al., 2021). We therefore hypothesised that SARS-CoV-2 may induce VAIL via the activity of E. Expression of Strep-tagged SARS-CoV-2 E protein in GFP-LC3B HEK293T cells resulted in the relocalisation GFP-LC3B to E-positive compartments (Figure 3A), while expression of mCherry had no effect on the distribution of LC3. This indicates that E is sufficient to induce LC3 lipidation.

**Figure 3:**
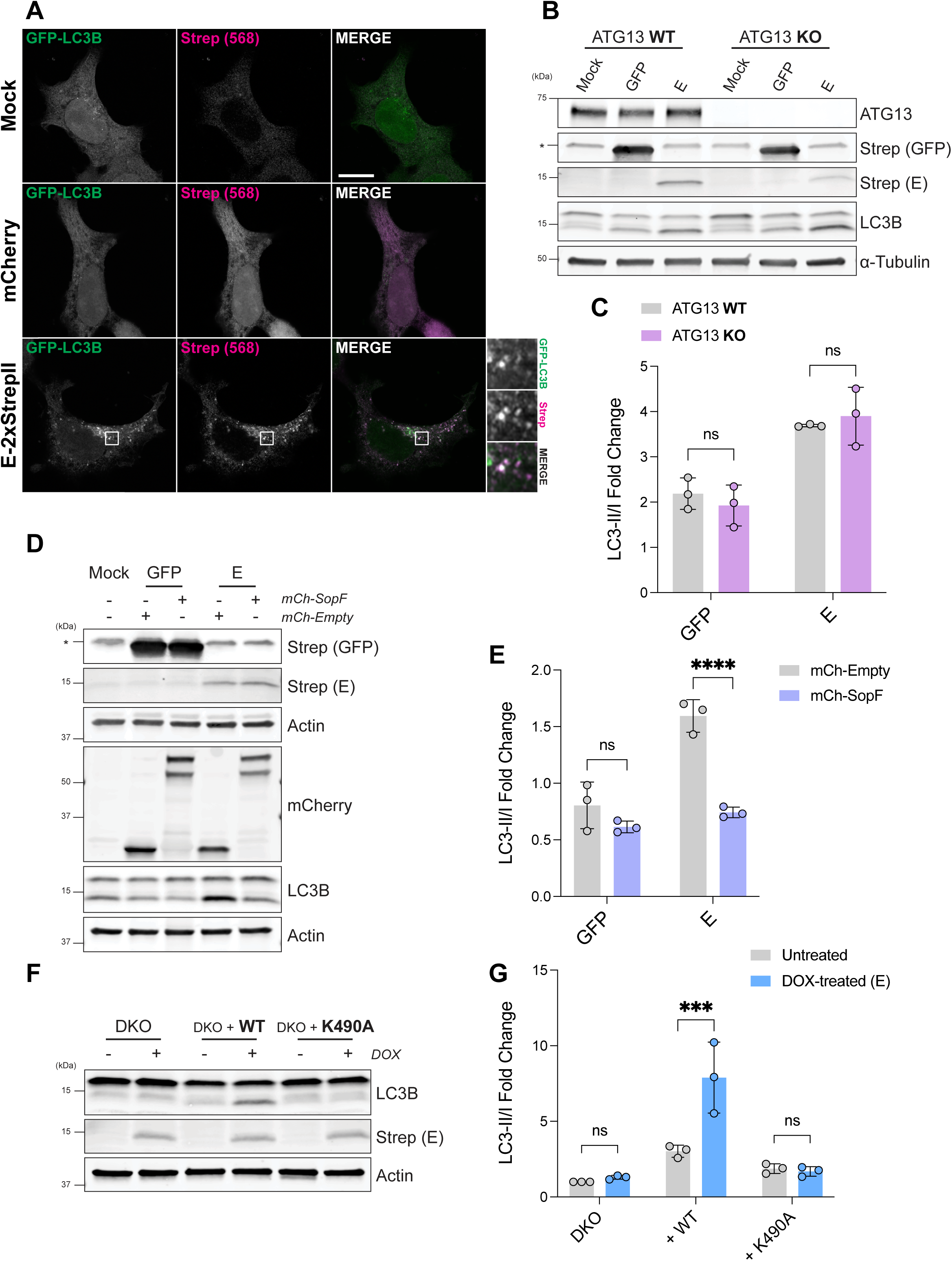
SARS-CoV-2 envelope induces VAIL A. Representative images of HEK293T cells expressing EGFP-LC3B (in green) that were transfected with E-2xStrep or mCherry, fixed after 24 h and stained with Strep (in magenta). Scale bar, 10 µm. B. Western blot analysis of ATG13 KO or WT HEK293T cells lysed 24 h after transfection of GFP-2xStrep or E-2xStrep. C. Quantification of LC3B-II/I ratios from B, relative to WT mock samples. Bars show mean ± SD; n=3. Two-way ANOVA with Šídák’s multiple comparisons. D. Western blot analysis of HEK293T cells lysed 48 h after co-transfections of EGFP-2xStrep or E-2xStrep with either mCherry-Empty or mCherry-SopF. E. Quantification of LC3B-II/I ratios from D, relative to mock samples. Bars show mean ± SD; n=3. ****: P ≤ 0.0001. Two-way ANOVA with Šídák’s multiple comparisons. F. ATG13/ATG16L1 double KO HEK293 cells (DKO) reconstituted with either WT or K490A FLAG-S-ATG16L1 were transduced with TetON E-2xStrep. Following doxycycline induction (10 μg/mL; 18h), cells were lysed and analysed by western blot. All cells were treated with MG132 (10 μM) for 2h prior to lysis to stabilise E-2xStrep. G. Quantification of LC3B-II/I ratios from F. Bars show mean ± SD; n=3. ***: P ≤ 0.001. Two-way ANOVA with Šídák’s multiple comparisons.

To assess whether LC3 lipidation triggered by E is driven by VAIL or canonical autophagy, we obtained ATG13 KO HEK293T cells (Kannangara et al., 2021). We confirmed that ATG13 KO cells treated with Torin 1 and bafilomycin A1, which in WT cells should result in the accumulation of autophagosome-conjugated LC3 (Yamamoto et al., 1998), showed no increase in LC3 lipidation (Figure S2A). Despite the deficiency in canonical autophagy, we observed no difference in E-induced LC3B lipidation in ATG13 KO cells relative to WT control cells (Figures 3B and 3C). Pharmacological inhibition of the Vps34 complex, which disrupts autophagy (Kim et al., 2013), had no impact on LC3 lipidation in E-expressing cells (Figures S2B and S2C). Together, these results suggest that the LC3 lipidation driven by SARS-CoV-2 E is independent of the canonical autophagy pathway.

We next examined the effect of SopF on the ability of E to induce LC3 lipidation. Co-expression of E with mCherry-Empty vector resulted in a clear increase in LC3B lipidation, as expected (Figures 3D and 3E). However, this LC3B lipidation was inhibited when E was co-expressed with mCherry-SopF, suggesting that E triggers LC3 lipidation via the VAIL pathway (Figures 3D and 3E). To further verify that E induces VAIL, we examined the effect of the K490A substitution in the WD40 domain of ATG16L1, which is known to disrupt V-ATPase-mediated ATG16L1 recruitment but not autophagy (Fletcher et al., 2018, Ulferts et al., 2021). ATG13/ATG16L1 double KO HEK293 cells reconstituted with either WT or K490A ATG16L1 were transduced with E-2xStrep under the control of a doxycycline-inducible promoter (TetON). We confirmed that cells expressing K490A ATG16L1 were deficient in VAIL activity following treatment with the known VAIL stimulus monensin (Figure S2D) (Florey et al., 2015, Fletcher et al., 2018). Induction of E expression with doxycycline had no effect on LC3B lipidation in ATG13/ATG16L1 double KO cells, as anticipated (Figures 3F and 3G). Lipidation in response to E was rescued by expression of WT ATG16L1, but not K490A ATG16L1, further demonstrating that the response to E is driven by the V-ATPase-ATG16L1 axis (Figures 3F and 3G). Additionally, as VAIL is sensitive to the V-ATPase inhibitor bafilomycin A1, we tested the effect of bafilomycin A1 on E-induced lipidation using ATG13 KO cells (to exclude the effects of bafilomycin on canonical autophagy) (Florey et al., 2015). Consistent with the induction of VAIL, lipidation in response to E expression was reduced by bafilomycin treatment (Figures S2E and S2F). Collectively these results show that SARS-CoV-2 E protein stimulates VAIL.

### SARS-CoV-2 envelope ion channel activity is required for VAIL during infection

VAIL is triggered by the disruption of intracellular proton gradients at acidic organelles (Durgan and Florey, 2022). Consistent with E inducing VAIL at physiologically acidified compartments, we observed co-localisation of SARS-CoV-2 E and GFP-LC3B at structures that were positive for the V-ATPase subunit V1D (Figure 4A). SARS-CoV-2 E is a 75 amino acid protein containing a C-terminal domain, a transmembrane channel-forming domain and an N-terminal domain (Figure 4B) (Mandala et al., 2020). Previous studies have shown that the neutralisation of organelle pH mediated by E can be inhibited by mutation of critical ion channel-lining residues; in particular, the N15A mutation (or equivalent) has been shown to disrupt the pH modulatory properties of E in multiple coronaviruses, including SARS-CoV-2 (Miura et al., 2023, Torres et al., 2007, Ruch and Machamer, 2012, Webb et al., 2022). To test whether the ion channel activity of E underlies the induction of VAIL, E^WT^ and ion channel-deficient E^N15A^ were expressed in HEK293T cells (Figure 4C). While E^WT^ expression was associated with a significant increase in LC3B lipidation versus GFP only controls, no significant increase in LC3B lipidation was observed with E^N15A^ (Figures 4C and 4D). Further confirming the importance of E ion channel activity for LC3B lipidation, E^N15A^ failed to induce a relocalisation of GFP-LC3B to E-containing structures within transfected cells (Figure 4E). Importantly, E^N15A^ was not defective in localisation to acidified compartments, as E^N15A^ was observed at TGN46-positive *trans*-Golgi compartments and LAMP1-positive compartments; only E^WT^ was able to induce the targeting of GFP-LC3B to these structures (Figures S3A and S3B).

**Figure 4:**
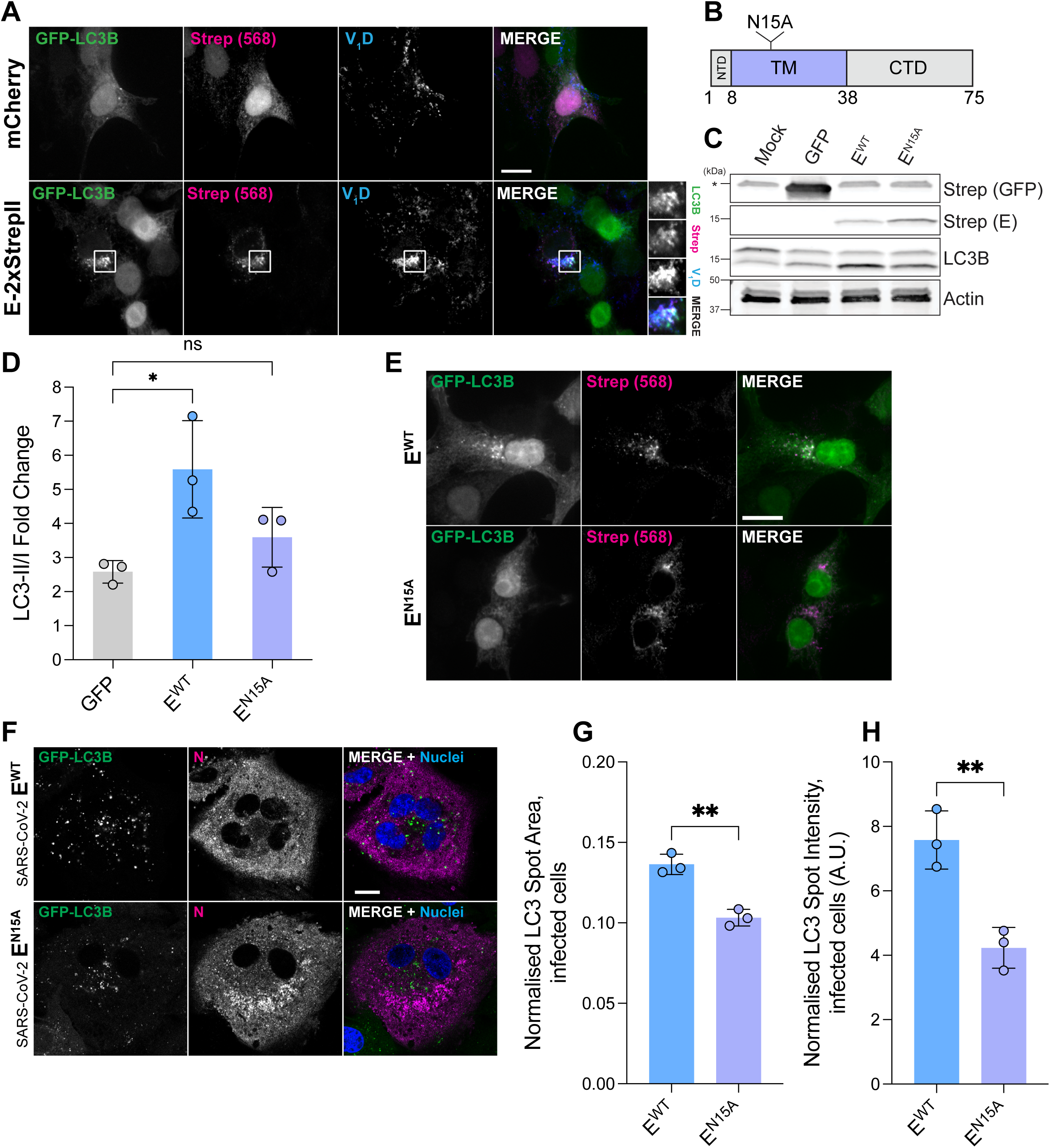
SARS-CoV-2 envelope ion channel activity is required for VAIL A. Representative images of HEK293T cells expressing EGFP-LC3B (in green) that were transfected with E-2xStrep or mCherry. Cells were fixed after 24 h and stained for V1D (in blue) and Strep (in magenta). Scale bar, 10 µm. B. Schematic of SARS-CoV-2 envelope. NTD: N-terminal domain; TM: transmembrane domain; CTD: C-terminal domain. C. Western blot analysis of HEK293T cells lysed 24 h after transfection with EGFP-2xStrep, E^WT^-2xStrep or E^N15A^-2xStrep. D. Quantification of LC3B-II/I ratios from C. Bars show mean ± SD; n=3. *: P ≤ 0.05. One-way ANOVA with Dunnett’s multiple comparisons. E. Representative images of HEK293T cells expressing EGFP-LC3B (in green) that were transfected with E^WT^-2xStrep or E^N15A^-2xStrep. Cells were fixed after 24 h and stained for Strep (in magenta). Scale bar, 10 µm. F. Representative images of A549 cells expressing ACE2/TMPRSS2 and EGFP-LC3B (in green). Indicated cells were infected with SARS-CoV-2 E^WT^ or SARS-CoV-2 E^N15A^ (MOI 2), fixed at 24 h post infection (hpi) and stained with SARS-CoV-2 nucleocapsid (N; in magenta) and NucBlue (nuclei; in blue). Scale bar, 10 µm. G. Quantification of EGFP-LC3B relocalisation in A549 cells expressing ACE2/TMPRSS2, EGFP-LC3B infected with either SARS-CoV-2 E^WT^ or SARS-CoV-2 E^N15A^ (MOI 2). Cells were fixed at 24 h post infection (hpi), stained with SARS-CoV-2 nucleocapsid and analysed using automated high-content imaging. The total area of EGFP-LC3B spots within infected cells was normalised to the total nucleocapsid-positive area in each well. Bars show mean ± SD; n=3. **: P ≤ 0.01. Unpaired *t*-test. H. Further quantification from G. The total intensity of EGFP-LC3B spots within infected cells was normalised to the total nucleocapsid-positive area in each well. Bars show mean ± SD; n=3. **: P ≤ 0.01. Unpaired *t*-test.

Having established the importance of E ion channel activity for VAIL induction in E-expressing cells, we aimed to investigate its contribution to LC3B lipidation in the context of whole virus infection. To do this, we generated SARS-CoV-2 virus containing the E^N15A^ mutation (on a hCoV-19/Wuhan/WIV04/2019 background) using a reverse genetics system (Figure S3C) (Zhou et al., 2022, Thi Nhu Thao et al., 2020). SARS-CoV-2 E^N15A^ virus was able to successfully infect A549 ACE2/TMPRSS2 cells, as determined by western blot and imaging, albeit to a reduced extent versus E^WT^ virus (Figures S3D and S3E). To account for the differences in infection efficiency, GFP-LC3B relocalisation was assessed in infected cells by fluorescence microscopy (Figure 4F). This approach revealed a partial reduction in GFP-LC3B punctum formation within cells infected with SARS-CoV-2 E^N15A^ relative to E^WT^, suggesting that E ion channel activity contributes directly to LC3 lipidation during infection (Figure 4F). This reduction in LC3 relocalisation was confirmed by automated imaging and analysis of A549 ACE2/TMPRSS2 GFP-LC3B cells infected with SARS-CoV-2 E^WT^ and E^N15A^ virus, both in terms of the area and intensity of LC3B puncta in infected cells (Figures 4G and 4H). Together, these data show that SARS-CoV-2 E ion channel activity is necessary for VAIL activation and contributes directly to LC3 lipidation during SARS-CoV-2 infection.

## Discussion

VAIL has been shown to be activated by a wide range of intracellular pathogens in response to infection-induced damage and perturbation of organelle pH (Biering et al., 2017, Beale et al., 2014, Ulferts et al., 2021, Wang et al., 2021). In this work, we find that SARS-CoV-2 stimulates VAIL, adding to the growing list of infectious agents that evoke this pathway. The initiation of VAIL during SARS-CoV-2 infection can be attributed, at least in part, to the ion channel activity of the viral envelope (E) protein. This activation of VAIL by a viral proton-conducting ion channel is reminiscent of processes that occur during other infections, particularly influenza A virus (IAV) (Fletcher et al., 2018). IAV induces VAIL via the proton channel activity of the viral ion channel M2, which shares a number of structural and functional properties with E (Schnell and Chou, 2008, Mandala et al., 2020, Breitinger et al., 2022). Both IAV- and SARS-CoV-2-induced VAIL play roles in the viral life cycle: in IAV infection, VAIL contributes virion stability and morphology (Beale et al., 2014, Figueras-Novoa et al., 2025); in SARS-CoV-2, the activation of VAIL by SARS-CoV-2 is required for efficient replication in cell culture. While some distinctions between VAIL observed in these two infections do exist, the similarities suggest common functions for VAIL during viral infection.

Our results are consistent with many previous studies which have described the induction of LC3 lipidation during SARS-CoV-2 infection (Gordon et al., 2020, Gassen et al., 2021, Miao et al., 2021, Shang et al., 2021, Qu et al., 2021, Gorshkov et al., 2021). However, these reports have focused largely on viral modulation of the canonical autophagy pathway. In this study, we employed established genetic tools to disentangle VAIL from autophagic processes, revealing new autophagy-independent functions of LC3 lipidation during SARS-CoV-2 infection. Inhibition of VAIL produced only a partial inhibition of LC3 lipidation during infection, indicating that previously reported alterations in autophagic flux likely occur alongside the activation of the V-ATPase-ATG16L1 axis.

The precise role of VAIL during SARS-CoV-2 infection remains an open question. Our findings suggest that VAIL plays a role in the later stages of the viral life cycle, given the minimal impact of SopF on pseudovirus entry and SARS-CoV-2 infection levels at early time points. SARS-CoV-2 and other betacoronaviruses have been described to exploit lysosomal compartments for egress (Pearson et al., 2024, Ghosh et al., 2020). Interestingly, multiple reports have shown that VAIL is able to promote lysosome biogenesis through activation of Transcription Factor EB (TFEB)-dependent transcription of lysosomal genes (Nakamura et al., 2020, Goodwin et al., 2021). Additional roles of VAIL may also be relevant for virion assembly and/or release. In particular, VAIL has been suggested to promote membrane remodelling and lipid trafficking through recruitment of the ESCRT machinery (endosomal sorting complex required for transport) (Ogura et al., 2023) and the lipid transferase ATG2 (Cross et al., 2023), respectively. Both IAV and SARS-CoV-2 derive their membranes from the host (Klein et al., 2020, Huang et al., 2022) and might exploit VAIL to promote optimal envelope formation.

While many studies have described the restriction of intracellular infection by VAIL (Wang et al., 2021, Selleck et al., 2015, Mitchell et al., 2018, Xu et al., 2019, Campbell-Valois et al., 2015, Köster et al., 2017, Choi et al., 2014, Biering et al., 2017, Hwang et al., 2012), we found that VAIL plays a pro-viral role during cellular infection with SARS-CoV-2. This suggests that the precise effect of VAIL is contextual and related to the particular pathogen involved. This mirrors the relationship between autophagy and infection, with the pathway playing both antagonistic and beneficial roles depending on the infectious agent and even the stage of infection (Jassey and Jackson, 2024). Attempts to manipulate autophagic pathways for therapeutic effect must therefore take account of the multiple different roles the autophagy machinery can play.

## Methods

### Antibodies and reagents

Antibodies used in this study were mouse monoclonal anti-Strep-tag II (Abcam, ab184224, WB 1:1000, IF 1:250), N human CR3009 antibody (Francis Crick Institute, described in (Zeng et al., 2021), IF 1:1000), N rabbit monoclonal antibody (Sino biological, 40143-R019, WB 1:1000), rabbit monoclonal anti-ATP6V1D (V1D) (Abcam, ab157458, WB 1:1000, IF 1:250), mouse monoclonal anti-β-actin (Proteintech, 66009-1, WB 1:10000), mouse monoclonal anti-GAPDH (Abcam, ab8245, WB 1:2000-5000), rabbit monoclonal anti-ATG16L1 (Cell Signalling Technology, 8089, WB 1:1000), rabbit monoclonal anti-LC3B (Novus Biologicals, NBP2-46892, WB 1:1000), rat monoclonal anti-α-tubulin (BioRad; MCA77G, WB 1:5000), rabbit monoclonal anti-ATG13 (Cell Signalling Technology, 13468, WB 1:1000), sheep polyclonal anti-TGN46 (BioRad, AHP500GT, IF 1:200), sheep polyclonal anti-LAMP1 (R&D Systems, AF4800, IF 1:200) and mouse monoclonal anti-mCherry (Abcam, ab18184, WB 1:1000).

Chemical reagents used in this study included monensin (Merck, M5273), bafilomycin A1 (Abcam, ab120497), Torin-1 (Selleckchem, S2827), Vps34 IN-1 compound 19 (Selleckchem, S8456), doxycycline (Sigma, D9891) and MG132 (Sigma, M7449).

### Cell culture

All cell lines were cultured in Dulbecco’s modified Eagle’s medium (DMEM; GIBCO Life Technologies) containing 10% Fetal Calf Serum (FCS), GlutaMax (GIBCO Life Technologies) and Penicillin-Streptomycin (GIBCO Life Technologies), unless otherwise indicated. Cells were grown at 37°C in 5% CO2. A549 and HEK293T cells were provided by Cell Services at the Francis Crick Institute. Vero E6 cells were from ATCC (catalogue code ATCC-CRL-1586). ATG13 KO HEK293T (with wild type parental controls) were a kind gift from J. Andersen from (Kannangara et al., 2021). ATG13/ATG16L1 double KO GFP-LC3B HEK293 cells (complemented with either WT or K490A FLAG-S-ATG16L1) were a kind gift from O. Florey.

### Plasmids

pLVX-EF1alpha-SARS-CoV-2-E-2xStrep-IRES-Puro (https://www.addgene.org/141385) and pLVX-EF1alpha-eGFP-2xStrep-IRES-Puro (https://www.addgene.org/141395) were gifts from N. Krogan (Gordon et al., 2020). pLVX-EF1alpha-SARS-CoV-2-E^N15A^-2xStrep-IRES-Puro was produced using the Q5 Site-Directed Mutagenesis Kit (New England BioLabs). pENTR-SARS-CoV-2-E-2xStrep was generated using the pENTR/D-TOPO Cloning Kit (Invitrogen). pInducer20-SARS-CoV-2-E-2xStrep-Neo (TetON E-2xStrep) was then produced by gateway cloning with LR clonase (Invitrogen), using pENTR-SARS-CoV-2-E-2xStrep and pInducer20-GWT-Neo. M5P-mCherry-SopF and M5P-mCherry-Empty control plasmid were described previously(Ulferts et al., 2021). pmCherry-N1, which was used as a transfection control in immunofluorescence experiments, was a kind gift from C. Crump. pWPI-IRES-Bla-Ak-ACE2-TMPRSS2 (https://www.addgene.org/154983) was a gift from S. Best. M4P-EGFP-LC3B and pOGP were kind gifts from F. Randow. pMD2.G (https://addgene.org/12259) and psPAX2 (https://addgene.org/12260) were gifts from D. Trono.

### Transient transfection

Cells were seeded in 24-, 12-, or 6-well plates and cultured for 24 h. Expression constructs were then mixed with PEI (polyethylenimine; 5 µg PEI per µg DNA) in Opti-MEM (GIBCO Life Technologies). After incubation at room temperature for 20 min, the DNA/PEI mixture was added dropwise to wells. Cells were then cultured for a further 24-48 h, as indicated in the relevant figures.

### Retrovirus and lentivirus generation

To produce Moloney Murine Leukaemia Virus (MMLV) retrovirus, transfer plasmids containing the insert of interest were co-transfected with pOGP and pMD2.G using PEI in HEK293T cells. Lentiviruses were produced in the same manner but with psPAX2 in place of pOGP. Cells were transduced with MMLV or lentivirus by spinoculation, at 500*g*, 1 h at room temperature, in the presence of 8 µg/mL polybrene. Transduced cells were selected using fluorescence-assisted cell sorting or 400 µg/mL G418 (Invivogen) as appropriate.

### siRNA knock-down

Cells were seeded in 6-well plates in fully complemented medium and incubated at 37°C in 5% CO2. After 48 h, 40 nM of siGENOME Non-Targeting Control siRNA Pool #1 (NT1; Horizon Discovery; D-001206-13-05), ON-TARGETplus SMARTPool human ATG16L1 siRNA (Horizon Discovery, L-021033-01-0005) or ON-TARGETplus SMARTPool human ATG13 siRNA (Horizon Discovery, L-020765-01-0005) was transfected using Lipofectamine RNAiMAX (Thermo), according to the manufacturer’s instructions. At 7 h post-transfection, medium in each well was replaced with fully complemented medium. After a further 48 h, a second round of siRNA transfection was performed, as above. Cells were infected with SARS-CoV-2 at 48 h after the second transfection, and lysed after the indicated length of infection.

### SDS-PAGE and western blot

Samples were lysed in ice-cold NP-40 buffer (0.5% NP-40, 25mM Tris-HCl (pH 7.5), 100 mM NaCl, 50 mM NaF) with fresh protease inhibitor cocktail (Sigma). Cell lysates were collected and clarified by centrifugation for 30 min at 4°C, 16,200*g*.

Protein concentration was determined using BCA assay (Pierce). Mini-PROTEAN®TGX gels (Bio-Rad) were used to separate proteins. Proteins were then transferred onto nitrocellulose membranes for western blot. Membranes were blocked with 5% dry milk powder in TBS with 0.1% Tween-20 for 30 min at room temperature.

Membranes were incubated with primary antibodies for 1 h at room temperature, or overnight at 4°C. After washing in TBS with 0.1% Tween-20, membranes were incubated with species-specific secondary antibodies coupled to IRdye 800CW or IRdye 680RD (LI-COR). Membranes were imaged using an Odyssey CLx scanner (LI-COR).

### Immunofluorescence and microscopy

Cells were grown on precision cover glasses (Marienfeld) or flat-bottom 96-well clear plates (PerkinElmer) coated with 0.001% poly-L-lysine (Sigma). After the desired treatment, cells were fixed with a final concentration of 4% formaldehyde in PBS for 20-60 min. Cells were washed with PBS and permeabilised with 0.2% Triton-X100 for 5 min. Permeabilised cells were blocked with 3% BSA in PBS for 30 min, prior to incubation with primary antibodies for 1 h (in 3% BSA/PBS). Samples were washed again with PBS and incubated for 45 min with species-specific secondary antibodies coupled to AlexaFluor 488, 568, or 647 (Thermo). Coverslips were mounted onto slides with ProLong^TM^ Glass Antifade Mountant (Invitrogen). For nuclear staining, coverslips were either mounted in ProLong^TM^ Glass Antifade Mountant with NucBlue Stain (Invitrogen), or incubated for 15 min with Hoechst 33342 (Invitrogen) prior to mounting. Images of stained coverslips were acquired using either a Zeiss LSM800 confocal microscope equipped with Airyscan (63x oil immersion lens) or a VisiTech iSIM microscope (100x oil immersion lens). Images were analysed and processed using Fiji (v.2.3.0/1.53) (Schindelin et al., 2012) and Adobe Photoshop 2023 (v.24.7.0).

### High throughput immunofluorescence quantification

Cells were cultured and infected on 96-well CellCarrier plates (PerkinElmer, 6005430). Plates were then scanned using Opera Phenix plus (PerkinElmer) after staining. The images were acquired utilizing a 40x_water_NA1.1 lens with confocal of 8 planes Z-stacks spanning from −1 to 2.5 µM. The process involved the utilization of excitation lasers at wavelengths of 405, 488, 561, and 647 nm. N and GFP-LC3B levels were analysed using Harmony 5.0.

### SARS-CoV-2 virus production, plaque assays and infection

To generate and propagate SARS-CoV-2 viral stocks, Vero V1 cells were seeded 48 h prior to infection, to achieve 70% confluency, in fully complemented DMEM (10% FCS; Pen/Strep; 37°C, 5% CO2). Cells were washed twice with serum-free DMEM, before virus diluted in serum-free media was added for 1 h before being replaced with serum-free media containing 1µg/mL TPCK-treated Trypsin. Supernatant was harvested once CPE was observed or 72 h post-infection. Viral supernatants were clarified by centrifugation at 2,000rpm for 15 min, and stored at −80°C until use.

To carry out plaque assays, Vero E6 cells were seeded in 12-well plates 48 h prior to infection, to achieve 100% confluency, in fully complemented DMEM. Monolayers were washed twice with serum-free media. Serial dilutions of viral inoculum were added to cell monolayers and incubated for 1 h. Inoculum was replaced with Avicel overlay (1.2% Avicel; 1X MEM; 0.01% DEAE Dextran; 1X MEM non-essential amino acids; 1µg/mL TPCK Trypsin) and plates were incubated for a further 72 h. Cells were then fixed with 4% paraformaldehyde and stained with 0.2% toluidine blue diluted in PBS.

To infect cells, wells were washed with serum free media, and incubated with inoculum containing virus (corresponding to the multiplicity of infection (MOI) given in figure legends) for 1 h. Inoculum was then removed, and fully complemented media was added. Cells were incubated for the period of time indicated in the legends of relevant figures before fixation or lysis.

### Reverse genetics

The E N15A virus and the control Wuhan virus were generated by reverse genetics using a previously described yeast-based system (Zhou et al., 2022, Thi Nhu Thao et al., 2020). Briefly, a series of overlapping cDNA fragments that span the complete SARS-CoV-2 Wuhan sequence (GenBank:MN908947.3) were chemically synthesized and cloned into pUC57-Kan (Biobasic Canada Inc.) The E N15A mutation was introduced into the fragment containing the E gene by site-directed mutagenesis and verified by Sanger sequencing. The SARS-CoV-2 fragments together with fragments containing the T7 RNA polymerase sequence followed by the 5’ end of the virus and the 3’ end of the virus followed by the hepatitis delta virus ribozyme and termination sequences from Vesicular Stomatitis Virus (VSV) and T7 bacteriophage were prepared either by PCR amplification or restriction digestion and gel purification. The viral genome was assembled from the fragments into the vector creating a full-length cDNA template using transformation-associated recombination (TAR) in TYC1 yeast. Full-length viral RNA transcripts were generated from the cDNA template *in vitro* using the Ribomax T7 RNA transcription kit and Ribo m7G Cap Analogue (Promega). The capped RNA transcripts were transfected into BHK-hACE2-N cells which stably express both the human ACE2 gene and the SARS-Cov-2 N gene to rescue virus. The rescued virus was passaged in Vero.E6 cells to generate a P1 stock and the genomes were sequenced using Oxford Nanopore as previously described (da Silva Filipe et al., 2021).

### RT-qPCR

RNA was isolated using RNAeasy extraction Kit Qiagen according to the manufacturer’s instructions. cDNA was synthesised using SuperSCRIPT-II reverse transcriptase (Invitrogen, 18064014) and PCR nucleotide mix (Promega, C1141) according to the manufacturer’s instructions. qPCR was performed using taq PCR and cycler using the following primers: SARS-CoV-2 N: CACATTGGCACCCGCAATC and GAGGAACGAGAAGAGGCTTG; SARS-CoV-2 RdRp: GTGARATGGTCATGTGTGGCGG and CARATGTTAAASACACTATTAGCATA; GAPDH: GGAGCGAGATCCCTCCAAAAT and GGCTGTTGTCATACTTCTCATGG.

### Pseudovirus entry assay

Lentiviral particles pseudotyped with SARS-CoV-2 Spike were produced by co-transfection of HEK293T cells with plasmids encoding Spike glycoproteins together with a plasmid encoding the SIVmac Gag-Pol polyprotein and a plasmid expressing an HIV-2 backbone with a GFP encoding gene, using GeneJuice (EMD Millipore).

Virus-containing supernatants were collected 48 h post-transfection and stored at −80°C until further use. For entry assays, lentiviral pseudotypes were added to cells seeded in 96-well plates (3,000 cells/well). Polybrene (4 µg/ml, Sigma Aldrich) was also added to the cells and plates were spun at 315*g* for 45 min. The percentage of transduced (GFP+) cells was assessed by flow cytometry 72 h later.

## Statistical analysis

Statistical analysis was performed in GraphPad Prism 9 (v.9.4.0). Statistical tests, number of replicates, and statistical significance are reported in specific experiment figure legends.

## Acknowledgements

The authors thank past and present members of the Beale for helpful comments and suggestions. The authors thank Svend Kjaer, from the Structural Biology STP, and High Throughput Screening, Cell Science, and Advanced Light Microscopy core facilities (Francis Crick Institute) for support. The authors also thank Dr Joshua L Andersen, Brigham Young University, USA for the 293T ATG13 KO cells. CFN, LT, EM, RTM, LA, MJ, MW, BM, KN, GK, RH, MH, RU, and RB were supported by The Francis Crick Institute, which receives its core funding from Cancer Research UK (CC2087, CC2088, CC1114), the UK Medical Research Council (CC2087, CC2088, CC1114) and the Wellcome Trust (CC2087, CC2088, CC1114). GDL, WF, VMC, NU, and AHP receive funding from the G2P-UK National Virology Consortium funded by the MRC (MR/W005611/1) and MRC grants (MC_UU12014/2 and MC_UU_00034/9) and the Wellcome Trust (206369/Z/17/2). For the purpose of Open Access, the author has applied a CC BY public copyright licence to any Author Accepted Manuscript version arising from this submission.

## Supplementary Figures

**Figure S1:** Supplementary data supporting the role of VAIL in SARS-CoV-2 replication A. Mean mCherry intensity within infected A549 cells expressing ACE2/TMPRSS2 and either mCherry-Empty or mCherry-SopF. Cells were infected with SARS-CoV-2 at the indicated MOI, fixed at 24 h post infection (hpi), stained with SARS-CoV-2 nucleocapsid and analysed using automated high-content imaging. Bars show mean ± SD; n=4. Two-way ANOVA with Šídák’s multiple comparisons. B. Mean nucleocapsid (N) intensity within infected A549 cells expressing ACE2/TMPRSS2 and either mCherry-Empty or mCherry-SopF. Cells were infected with SARS-CoV-2 at the indicated MOI, fixed at 24 h post infection (hpi), stained with SARS-CoV-2 nucleocapsid and analysed using automated high-content imaging. Bars show mean ± SD; n=4. Two-way ANOVA with Šídák’s multiple comparisons. C. Titres (in plaque forming units (PFU) per mL) of SARS-CoV-2 virus passaged in A549 cells expressing ACE2/TMPRSS2 and either mCherry-Empty or mCherry-SopF. Indicated cells were infected at an MOI of 5 and the supernatant was collected at 24 h post infection. Titres were determined by Vero E6 plaque assay.

**Figure S2:** Supplementary data supporting the induction of VAIL by SARS-CoV-2 envelope. A. Western blot analysis of ATG13 KO or WT HEK293T cells lysed after 4 h treatment with 250 nM Torin-1 and 100 nM bafilomycin A1 (Baf), or DMSO only. B. Western blot analysis of HEK293T cells lysed 24 h after transfection of EGFP-2xStrep or E-2xStrep. 1 µM Vps34 IN-1 was added 4 h prior to lysis, as indicated. C. Quantification of LC3B-II/I ratios from E. Bars show mean ± SD; n=3. **: P ≤ 0.01. Two-way ANOVA with Šídák’s multiple comparisons. D. Western blot analysis of ATG13/ATG16L1 double KO HEK293 cells (DKO) reconstituted with either WT or K490A FLAG-S-ATG16L1 and treated with 100 µM monensin for 1 h. E. Western blot analysis of ATG13 KO HEK293T cells lysed 24 h after transfection of EGFP-2xStrep or E-2xStrep. 100 nM bafilomycin A1 was added 4 h prior to lysis, as indicated. F. Quantification of LC3B-II/I ratios from C. Bars show mean ± SD; n=3. **: P ≤ 0.01. Two-way ANOVA with Šídák’s multiple comparisons.

**Figure S3:** Supplementary data supporting the role of SARS-CoV-2 envelope ion channel activity in VAIL. A. Representative images of HEK293T cells expressing EGFP-LC3B (in green) that were transfected with either mCherry, E^WT^-2xStrep, or E^N15A^-2xStrep. Cells were fixed after 24 h and stained for TGN46 (in blue) and Strep (in magenta). Scale bar, 10 µm. Representative images of EGFP-LC3B HEK293T cells that were transfected with either mCherry, E^WT^-2xStrep, or E^N15A^-2xStrep. Cells were fixed after 24 h and stained for LAMP1 (in blue) and Strep (in magenta). Scale bar, 10 µm. Sequencing of the SARS-CoV-2 E^N15A^ mutant virus aligned to the hCoV-19/Wuhan/WIV04/2019 sequence. The start of the envelope reading frame is shown, with the WT amino acid translation shown below. Two nucleotides, highlighted, were changed to encode the N15A mutation. Western blot of A549 cells expressing ACE2/TMPRSS2. Cells were infected with either SARS-CoV-2 E^WT^ or SARS-CoV-2 E^N15A^ (MOI 2) and lysed 24 h post infection (hpi). Percentage of infected (nucleocapsid-positive) A549 cells expressing ACE2/TMPRSS2. Cells were infected with either SARS-CoV-2 E^WT^ or SARS-CoV-2 E^N15A^ (MOI 2), fixed at 24 h post infection (hpi), stained with SARS-CoV-2 nucleocapsid and analysed using automated high-content imaging. Bars show mean ± SD; n=3. ***: P ≤ 0.001. One-way ANOVA with Tukey’s multiple comparisons.

## REFERENCES

Beale, R., Wise, H., Stuart, A., Ravenhill, B. J., Digard, P. & Randow, F. 2014. A LC3-interacting motif in the influenza A virus M2 protein is required to subvert autophagy and maintain virion stability. Cell Host Microbe, 15, 239–47.

Biering, S. B., Choi, J., Halstrom, R. A., Brown, H. M., Beatty, W. L., Lee, S., Mccune, B. T., Dominici, E., Williams, L. E., Orchard, R. C., Wilen, C. B., Yamamoto, M., Coers, J., Taylor, G. A. & Hwang, S. 2017. Viral Replication Complexes Are Targeted by LC3-Guided Interferon-Inducible GTPases. Cell Host Microbe, 22, 74–85.e7.

Breitinger, U., Farag, N. S., Sticht, H. & Breitinger, H.-G. 2022. Viroporins: Structure, function, and their role in the life cycle of SARS-CoV-2. The International Journal of Biochemistry & Cell Biology, 145, 106185.

Cabrera-Garcia, D., Bekdash, R., Abbott, G. W., Yazawa, M. & Harrison, N. L. 2021. The envelope protein of SARS-CoV-2 increases intra-Golgi pH and forms a cation channel that is regulated by pH. J Physiol, 599, 2851–2868.

Campbell-Valois, F. X., Sachse, M., Sansonetti, P. J. & Parsot, C. 2015. Escape of Actively Secreting Shigella flexneri from ATG8/LC3-Positive Vacuoles Formed during Cell-To-Cell Spread Is Facilitated by IcsB and VirA. mBio, 6, e02567–14.

Choi, J., Park, S., Biering, S. B., Selleck, E., Liu, C. Y., Zhang, X., Fujita, N., Saitoh, T., Akira, S., Yoshimori, T., Sibley, L. D., Hwang, S. & Virgin, H. W. 2014. The parasitophorous vacuole membrane of Toxoplasma gondii is targeted for disruption by ubiquitin-like conjugation systems of autophagy. Immunity, 40, 924–35.

Ciampor, F., Bayley, P. M., Nermut, M. V., Hirst, E. M., Sugrue, R. J. & Hay, A. J. 1992. Evidence that the amantadine-induced, M2-mediated conversion of influenza A virus hemagglutinin to the low pH conformation occurs in an acidic trans Golgi compartment. Virology, 188, 14–24.

Cross, J., Durgan, J., Mcewan, D. G., Tayler, M., Ryan, K. M. & Florey, O. 2023. Lysosome damage triggers direct ATG8 conjugation and ATG2 engagement via non-canonical autophagy. J Cell Biol, 222.

Da Silva Filipe, A., Shepherd, J. G., Williams, T., Hughes, J., Aranday-Cortes, E., Asamaphan, P., Ashraf, S., Balcazar, C., Brunker, K., Campbell, A., Carmichael, S., Davis, C., Dewar, R., Gallagher, M. D., Gunson, R., Hill, V., Ho, A., Jackson, B., James, E., Jesudason, N., Johnson, N., Mcwilliam Leitch, E. C., Li, K., Maclean, A., Mair, D., Mcallister, D. A., Mccrone, J. T., Mcdonald, S. E., Mchugh, M. P., Morris, A. K., Nichols, J., Niebel, M., Nomikou, K., Orton, R. J., O’toole, Á., Palmarini, M., Parcell, B. J., Parr, Y. A., Rambaut, A., Rooke, S., Shaaban, S., Shah, R., Singer, J. B., Smollett, K., Starinskij, I., Tong, L., Sreenu, V. B., Wastnedge, E., Holden, M. T. G., Robertson, D. L., Templeton, K. & Thomson, E. C. 2021. Genomic epidemiology reveals multiple introductions of SARS-CoV-2 from mainland Europe into Scotland. Nat Microbiol, 6, 112–122.

Durgan, J. & Florey, O. 2022. Many roads lead to CASM: Diverse stimuli of noncanonical autophagy share a unifying molecular mechanism. Sci Adv, 8, eabo1274.

Figueras-Novoa, C., Akutsu, M., Murata, D., Weston, A., Jiang, M., Montaner, B., Dubois, C., Shenoy, A. & Beale, R. 2025. Caspase cleavage of influenza A virus M2 disrupts M2-LC3 interaction and regulates virion production. EMBO reports, 26, 1768–1791-1791.

Figueras-Novoa, C., Timimi, L., Marcassa, E., Ulferts, R. & Beale, R. 2024. Conjugation of ATG8s to single membranes at a glance. Journal of Cell Science, 137, jcs261031.

Fischer, T. D., Wang, C., Padman, B. S., Lazarou, M. & Youle, R. J. 2020. STING induces LC3B lipidation onto single-membrane vesicles via the V-ATPase and ATG16L1-WD40 domain. Journal of Cell Biology, 219, e202009128.

Fletcher, K., Ulferts, R., Jacquin, E., Veith, T., Gammoh, N., Arasteh, J. M., Mayer, U., Carding, S. R., Wileman, T., Beale, R. & Florey, O. 2018. The WD40 domain of ATG16L1 is required for its non-canonical role in lipidation of LC3 at single membranes. The EMBO journal, 37, e97840.

Florey, O., Gammoh, N., Kim, S. E., Jiang, X. & Overholtzer, M. 2015. V-ATPase and osmotic imbalances activate endolysosomal LC3 lipidation. Autophagy, 11, 88–99.

Florey, O., Kim, S. E., Sandoval, C. P., Haynes, C. M. & Overholtzer, M. 2011. Autophagy machinery mediates macroendocytic processing and entotic cell death by targeting single membranes. Nat Cell Biol, 13, 1335–43.

Gassen, N. C., Papies, J., Bajaj, T., Emanuel, J., Dethloff, F., Chua, R. L., Trimpert, J., Heinemann, N., Niemeyer, C., Weege, F., Hönzke, K., Aschman, T., Heinz, D. E., Weckmann, K., Ebert, T., Zellner, A., Lennarz, M., Wyler, E., Schroeder, S., Richter, A., Niemeyer, D., Hoffmann, K., Meyer, T. F., Heppner, F. L., Corman, V. M., Landthaler, M., Hocke, A. C., Morkel, M., Osterrieder, N., Conrad, C., Eils, R., Radbruch, H., Giavalisco, P., Drosten, C. & Müller, M. A. 2021. SARS-CoV-2-mediated dysregulation of metabolism and autophagy uncovers host-targeting antivirals. Nat Commun, 12, 3818.

Ghosh, S., Dellibovi-Ragheb, T. A., Kerviel, A., Pak, E., Qiu, Q., Fisher, M., Takvorian, P. M., Bleck, C., Hsu, V. W., Fehr, A. R., Perlman, S., Achar, S. R., Straus, M. R., Whittaker, G. R., De Haan, C. A. M., Kehrl, J., Altan-Bonnet, G. & Altan-Bonnet, N. 2020. β-Coronaviruses Use Lysosomes for Egress Instead of the Biosynthetic Secretory Pathway. Cell, 183, 1520–1535.e14.

Gluschko, A., Herb, M., Wiegmann, K., Krut, O., Neiss, W. F., Utermöhlen, O., Krönke, M. & Schramm, M. 2018. The β(2) Integrin Mac-1 Induces Protective LC3-Associated Phagocytosis of Listeria monocytogenes. Cell Host Microbe, 23, 324–337.e5.

Goodwin, J. M., Walkup, W. G. T., Hooper, K., Li, T., Kishi-Itakura, C., Ng, A., Lehmberg, T., Jha, A., Kommineni, S., Fletcher, K., Garcia-Fortanet, J., Fan, Y., Tang, Q., Wei, M., Agrawal, A., Budhe, S. R., Rouduri, S. R., Baird, D., Saunders, J., Kiselar, J., Chance, M. R., Ballabio, A., Appleton, B. A., Brumell, J. H., Florey, O. & Murphy, L. O. 2021. GABARAP sequesters the FLCN-FNIP tumor suppressor complex to couple autophagy with lysosomal biogenesis. Sci Adv, 7, eabj2485.

Gordon, D. E., Jang, G. M., Bouhaddou, M., Xu, J., Obernier, K., White, K. M., O’meara, M. J., Rezelj, V. V., Guo, J. Z., Swaney, D. L., Tummino, T. A., Hüttenhain, R., Kaake, R. M., Richards, A. L., Tutuncuoglu, B., Foussard, H., Batra, J., Haas, K., Modak, M., Kim, M., Haas, P., Polacco, B. J., Braberg, H., Fabius, J. M., Eckhardt, M., Soucheray, M., Bennett, M. J., Cakir, M., Mcgregor, M. J., Li, Q., Meyer, B., Roesch, F., Vallet, T., Mac Kain, A., Miorin, L., Moreno, E., Naing, Z. Z. C., Zhou, Y., Peng, S., Shi, Y., Zhang, Z., Shen, W., Kirby, I. T., Melnyk, J. E., Chorba, J. S., Lou, K., Dai, S. A., Barrio-Hernandez, I., Memon, D., Hernandez-Armenta, C., Lyu, J., Mathy, C. J. P., Perica, T., Pilla, K. B., Ganesan, S. J., Saltzberg, D. J., Rakesh, R., Liu, X., Rosenthal, S. B., Calviello, L., Venkataramanan, S., Liboy-Lugo, J., Lin, Y., Huang, X.-P., Liu, Y., Wankowicz, S. A., Bohn, M., Safari, M., Ugur, F. S., Koh, C., Savar, N. S., Tran, Q. D., Shengjuler, D., Fletcher, S. J., O’neal, M. C., Cai, Y., Chang, J. C. J., Broadhurst, D. J., Klippsten, S., Sharp, P. P., Wenzell, N. A., Kuzuoglu-Ozturk, D., Wang, H.-Y., Trenker, R., Young, J. M., Cavero, D. A., Hiatt, J., Roth, T. L., Rathore, U., Subramanian, A., Noack, J., Hubert, M., Stroud, R. M., Frankel, A. D., Rosenberg, O. S., Verba, K. A., Agard, D. A., Ott, M., Emerman, M., Jura, N., et al. 2020. A SARS-CoV-2 protein interaction map reveals targets for drug repurposing. Nature, 583, 459–468.

Gorshkov, K., Chen, C. Z., Bostwick, R., Rasmussen, L., Tran, B. N., Cheng, Y. S., Xu, M., Pradhan, M., Henderson, M., Zhu, W., Oh, E., Susumu, K., Wolak, M., Shamim, K., Huang, W., Hu, X., Shen, M., Klumpp-Thomas, C., Itkin, Z., Shinn, P., Carlos De La Torre, J., Simeonov, A., Michael, S. G., Hall, M. D., Lo, D. C. & Zheng, W. 2021. The SARS-CoV-2 Cytopathic Effect Is Blocked by Lysosome Alkalizing Small Molecules. ACS Infect Dis, 7, 1389–1408.

Hooper, K. M., Jacquin, E., Li, T., Goodwin, J. M., Brumell, J. H., Durgan, J. & Florey, O. 2022. V-ATPase is a universal regulator of LC3-associated phagocytosis and non-canonical autophagy. J Cell Biol, 221.

Huang, Q. J., Song, K., Xu, C., Bolon, D. N. A., Wang, J. P., Finberg, R. W., Schiffer, C. A. & Somasundaran, M. 2022. Quantitative structural analysis of influenza virus by cryo-electron tomography and convolutional neural networks. Structure, 30, 777–786.e3.

Hwang, S., Maloney, N. S., Bruinsma, M. W., Goel, G., Duan, E., Zhang, L., Shrestha, B., Diamond, M. S., Dani, A., Sosnovtsev, S. V., Green, K. Y., Lopez-Otin, C., Xavier, R. J., Thackray, L. B. & Virgin, H. W. 2012. Nondegradative role of Atg5-Atg12/ Atg16L1 autophagy protein complex in antiviral activity of interferon gamma. Cell host & microbe, 11, 397–409.

Jackson, C. B., Farzan, M., Chen, B. & Choe, H. 2022. Mechanisms of SARS-CoV-2 entry into cells. Nature Reviews Molecular Cell Biology, 23, 3–20.

Jassey, A. & Jackson, W. T. 2024. Viruses and autophagy: bend, but don’t break. Nature Reviews Microbiology, 22, 309–321.

Jung, C. H., Jun, C. B., Ro, S. H., Kim, Y. M., Otto, N. M., Cao, J., Kundu, M. & Kim, D. H. 2009. ULK-Atg13-FIP200 complexes mediate mTOR signaling to the autophagy machinery. Mol Biol Cell, 20, 1992–2003.

Kannangara, A. R., Poole, D. M., Mcewan, C. M., Youngs, J. C., Weerasekara, V. K., Thornock, A. M., Lazaro, M. T., Balasooriya, E. R., Oh, L. M., Soderblom, E. J., Lee, J. J., Simmons, D. L. & Andersen, J. L. 2021. BioID reveals an ATG9A interaction with ATG13-ATG101 in the degradation of p62/SQSTM1-ubiquitin clusters. EMBO reports, 22, e51136.

Kim, J., Kim, Y. C., Fang, C., Russell, R. C., Kim, J. H., Fan, W., Liu, R., Zhong, Q. & Guan, K. L. 2013. Differential regulation of distinct Vps34 complexes by AMPK in nutrient stress and autophagy. Cell, 152, 290–303.

Klein, S., Cortese, M., Winter, S. L., Wachsmuth-Melm, M., Neufeldt, C. J., Cerikan, B., Stanifer, M. L., Boulant, S., Bartenschlager, R. & Chlanda, P. 2020. SARS-CoV-2 structure and replication characterized by in situ cryo-electron tomography. Nat Commun, 11, 5885.

Köster, S., Upadhyay, S., Chandra, P., Papavinasasundaram, K., Yang, G., Hassan, A., Grigsby, S. J., Mittal, E., Park, H. S., Jones, V., Hsu, F. F., Jackson, M., Sassetti, C. M. & Philips, J. A. 2017. Mycobacterium tuberculosis is protected from NADPH oxidase and LC3-associated phagocytosis by the LCP protein CpsA. Proc Natl Acad Sci U S A, 114, E8711–e8720.

Linders, P. T. A., Ioannidis, M., Ter Beest, M. & Van Den Bogaart, G. 2022. Fluorescence Lifetime Imaging of pH along the Secretory Pathway. ACS Chemical Biology, 17, 240–251.

Liu, B., Carlson, R. J., Pires, I. S., Gentili, M., Feng, E., Hellier, Q., Schwartz, M. A., Blainey, P. C., Irvine, D. J. & Hacohen, N. 2023. Human STING is a proton channel. Science, 381, 508–514.

Mandala, V. S., Mckay, M. J., Shcherbakov, A. A., Dregni, A. J., Kolocouris, A. & Hong, M. 2020. Structure and drug binding of the SARS-CoV-2 envelope protein transmembrane domain in lipid bilayers. Nat Struct Mol Biol, 27, 1202–1208.

Mercer, J., Schelhaas, M. & Helenius, A. 2010. Virus entry by endocytosis. Annu Rev Biochem, 79, 803–33.

Miao, G., Zhao, H., Li, Y., Ji, M., Chen, Y., Shi, Y., Bi, Y., Wang, P. & Zhang, H. 2021. ORF3a of the COVID-19 virus SARS-CoV-2 blocks HOPS complex-mediated assembly of the SNARE complex required for autolysosome formation. Dev Cell, 56, 427–442.e5.

Mitchell, G., Cheng, M. I., Chen, C., Nguyen, B. N., Whiteley, A. T., Kianian, S., Cox, J. S., Green, D. R., Mcdonald, K. L. & Portnoy, D. A. 2018. Listeria monocytogenes triggers noncanonical autophagy upon phagocytosis, but avoids subsequent growth-restricting xenophagy. Proceedings of the National Academy of Sciences, 115, E210–E217.

Miura, K., Suzuki, Y., Ishida, K., Arakawa, M., Wu, H., Fujioka, Y., Emi, A., Maeda, K., Hamajima, R., Nakano, T., Tenno, T., Hiroaki, H. & Morita, E. 2023. Distinct motifs in the E protein are required for SARS-CoV-2 virus particle formation and lysosomal deacidification in host cells. Journal of Virology, 97, e00426–23.

Nakamura, S., Shigeyama, S., Minami, S., Shima, T., Akayama, S., Matsuda, T., Esposito, A., Napolitano, G., Kuma, A., Namba-Hamano, T., Nakamura, J., Yamamoto, K., Sasai, M., Tokumura, A., Miyamoto, M., Oe, Y., Fujita, T., Terawaki, S., Takahashi, A., Hamasaki, M., Yamamoto, M., Okada, Y., Komatsu, M., Nagai, T., Takabatake, Y., Xu, H., Isaka, Y., Ballabio, A. & Yoshimori, T. 2020. LC3 lipidation is essential for TFEB activation during the lysosomal damage response to kidney injury. Nature Cell Biology, 22, 1252–1263.

Ogura, M., Kaminishi, T., Shima, T., Torigata, M., Bekku, N., Tabata, K., Minami, S., Nishino, K., Nezu, A., Hamasaki, M., Kosako, H., Yoshimori, T. & Nakamura, S. 2023. Microautophagy regulated by STK38 and GABARAPs is essential to repair lysosomes and prevent aging. EMBO Rep, 24, e57300.

Pearson, G. J., Mears, H. V., Broncel, M., Snijders, A. P., Bauer, D. L. V. & Carlton, J. G. 2024. ER-export and ARFRP1/AP-1-dependent delivery of SARS-CoV-2 Envelope to lysosomes controls late stages of viral replication. Sci Adv, 10, eadl5012.

Qu, Y., Wang, X., Zhu, Y., Wang, W., Wang, Y., Hu, G., Liu, C., Li, J., Ren, S., Xiao, M. Z. X., Liu, Z., Wang, C., Fu, J., Zhang, Y., Li, P., Zhang, R. & Liang, Q. 2021. ORF3a-Mediated Incomplete Autophagy Facilitates Severe Acute Respiratory Syndrome Coronavirus-2 Replication. Front Cell Dev Biol, 9, 716208.

Ren, Y., Li, C., Feng, L., Pan, W., Li, L., Wang, Q., Li, J., Li, N., Han, L., Zheng, X., Niu, X., Sun, C. & Chen, L. 2016. Proton Channel Activity of Influenza A Virus Matrix Protein 2 Contributes to Autophagy Arrest. J Virol, 90, 591–8.

Ruch, T. R. & Machamer, C. E. 2012. A single polar residue and distinct membrane topologies impact the function of the infectious bronchitis coronavirus E protein. PLoS Pathog, 8, e1002674.

Schindelin, J., Arganda-Carreras, I., Frise, E., Kaynig, V., Longair, M., Pietzsch, T., Preibisch, S., Rueden, C., Saalfeld, S., Schmid, B., Tinevez, J. Y., White, D. J., Hartenstein, V., Eliceiri, K., Tomancak, P. & Cardona, A. 2012. Fiji: an open-source platform for biological-image analysis. Nat Methods, 9, 676–82.

Schnell, J. R. & Chou, J. J. 2008. Structure and mechanism of the M2 proton channel of influenza A virus. Nature, 451, 591–5.

Selleck, E. M., Orchard, R. C., Lassen, K. G., Beatty, W. L., Xavier, R. J., Levine, B., Virgin, H. W. & Sibley, L. D. 2015. A Noncanonical Autophagy Pathway Restricts Toxoplasma gondii Growth in a Strain-Specific Manner in IFN-γ-Activated Human Cells. mBio, 6, e01157–15.

Shang, C., Zhuang, X., Zhang, H., Li, Y., Zhu, Y., Lu, J., Ge, C., Cong, J., Li, T., Li, N., Tian, M., Jin, N. & Li, X. 2021. Inhibition of Autophagy Suppresses SARS-CoV-2 Replication and Ameliorates Pneumonia in hACE2 Transgenic Mice and Xenografted Human Lung Tissues. J Virol, 95, e0153721.

Thi Nhu Thao, T., Labroussaa, F., Ebert, N., V’kovski, P., Stalder, H., Portmann, J., Kelly, J., Steiner, S., Holwerda, M., Kratzel, A., Gultom, M., Schmied, K., Laloli, L., Hüsser, L., Wider, M., Pfaender, S., Hirt, D., Cippà, V., Crespo-Pomar, S., Schröder, S., Muth, D., Niemeyer, D., Corman, V. M., Müller, M. A., Drosten, C., Dijkman, R., Jores, J. & Thiel, V. 2020. Rapid reconstruction of SARS-CoV-2 using a synthetic genomics platform. Nature, 582, 561–565.

Timimi, L., Wrobel, A. G., Chiduza, G. N., Maslen, S. L., Torres-Méndez, A., Montaner, B., Davis, C., Minckley, T., Hole, K. L., Serio, A., Devine, M. J., Skehel, J. M., Rubinstein, J. L., Schreiber, A. & Beale, R. 2024. The V-ATPase/ATG16L1 axis is controlled by the V(1)H subunit. Mol Cell, 84, 2966–2983.e9.

Torres, J., Maheswari, U., Parthasarathy, K., Ng, L., Liu, D. X. & Gong, X. 2007. Conductance and amantadine binding of a pore formed by a lysine-flanked transmembrane domain of SARS coronavirus envelope protein. Protein science: a publication of the Protein Society, 16, 2065–2071.

Ulferts, R., Marcassa, E., Timimi, L., Lee, L. C., Daley, A., Montaner, B., Turner, S. D., Florey, O., Baillie, J. K. & Beale, R. 2021. Subtractive CRISPR screen identifies the ATG16L1/vacuolar ATPase axis as required for non-canonical LC3 lipidation. Cell Rep, 37, 109899.

Veiga, V. C. & Cavalcanti, A. B. 2023. Age, host response, and mortality in COVID-19. Eur Respir J, 62.

Wang, Y., Sharma, P., Jefferson, M., Zhang, W., Bone, B., Kipar, A., Bitto, D., Coombes, J. L., Pearson, T., Man, A., Zhekova, A., Bao, Y., Tripp, R. A., Carding, S. R., Yamauchi, Y., Mayer, U., Powell, P. P., Stewart, J. P. & Wileman, T. 2021. Non-canonical autophagy functions of ATG16L1 in epithelial cells limit lethal infection by influenza A virus. Embo j, 40, e105543.

Webb, I., Keep, S., Littolff, K., Stuart, J., Freimanis, G., Britton, P., Davidson, A. D., Maier, H. J. & Bickerton, E. 2022. The Genetic Stability, Replication Kinetics and Cytopathogenicity of Recombinant Avian Coronaviruses with a T16A or an A26F Mutation within the E Protein Is Cell-Type Dependent. Viruses, 14.

Westerbeck, J. W. & Machamer, C. E. 2019. The Infectious Bronchitis Coronavirus Envelope Protein Alters Golgi pH To Protect the Spike Protein and Promote the Release of Infectious Virus. Journal of Virology, 93, e00015–19.

White, J. M. & Whittaker, G. R. 2016. Fusion of Enveloped Viruses in Endosomes. Traffic, 17, 593–614.

Xu, Y., Zhou, P., Cheng, S., Lu, Q., Nowak, K., Hopp, A. K., Li, L., Shi, X., Zhou, Z., Gao, W., Li, D., He, H., Liu, X., Ding, J., Hottiger, M. O. & Shao, F. 2019. A Bacterial Effector Reveals the V-ATPase-ATG16L1 Axis that Initiates Xenophagy. Cell, 178, 552–566.e20.

Yamamoto, A., Tagawa, Y., Yoshimori, T., Moriyama, Y., Masaki, R. & Tashiro, Y. 1998. Bafilomycin A1 prevents maturation of autophagic vacuoles by inhibiting fusion between autophagosomes and lysosomes in rat hepatoma cell line, H-4-II- E cells. Cell Struct Funct, 23, 33–42.

Yamauchi, Y. & Helenius, A. 2013. Virus entry at a glance. Journal of Cell Science, 126, 1289–1295.

Zeng, J., Weissmann, F., Bertolin, A. P., Posse, V., Canal, B., Ulferts, R., Wu, M., Harvey, R., Hussain, S., Milligan, J. C., Roustan, C., Borg, A., Mccoy, L., Drury, L. S., Kjaer, S., Mccauley, J., Howell, M., Beale, R. & Diffley, J. F. X. 2021. Identifying SARS-CoV-2 antiviral compounds by screening for small molecule inhibitors of nsp13 helicase. Biochem J, 478, 2405–2423.

Zhou, J., Peacock, T. P., Brown, J. C., Goldhill, D. H., Elrefaey, A. M. E., Penrice- Randal, R., Cowton, V. M., De Lorenzo, G., Furnon, W., Harvey, W. T., Kugathasan, R., Frise, R., Baillon, L., Lassaunière, R., Thakur, N., Gallo, G., Goldswain, H., Donovan-Banfield, I., Dong, X., Randle, N. P., Sweeney, F., Glynn, M. C., Quantrill, J. L., Mckay, P. F., Patel, A. H., Palmarini, M., Hiscox, J. A., Bailey, D. & Barclay, W. S. 2022. Mutations that adapt SARS-CoV-2 to mink or ferret do not increase fitness in the human airway. Cell Rep, 38, 110344.

Zhou, P., Yang, X. L., Wang, X. G., Hu, B., Zhang, L., Zhang, W., Si, H. R., Zhu, Y., Li, B., Huang, C. L., Chen, H. D., Chen, J., Luo, Y., Guo, H., Jiang, R. D., Liu, M. Q., Chen, Y., Shen, X. R., Wang, X., Zheng, X. S., Zhao, K., Chen, Q. J., Deng, F., Liu, L. L., Yan, B., Zhan, F. X., Wang, Y. Y., Xiao, G. F. & Shi, Z. L. 2020. A pneumonia outbreak associated with a new coronavirus of probable bat origin. Nature, 579, 270–273.

