## Supplementary figures and images for "SARS-CoV-2 envelope protein induces LC3 lipidation via the V-ATPase-ATG16L1 axis"

### Supplementary Figures

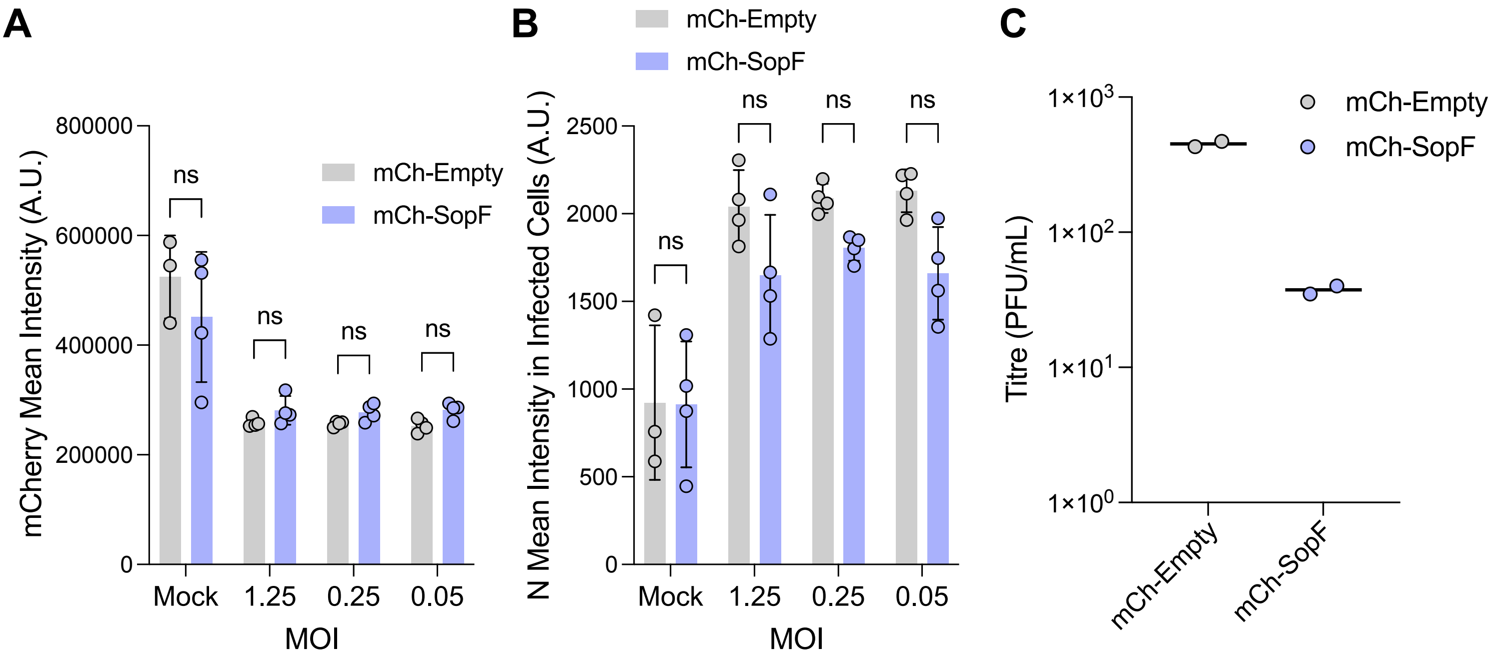

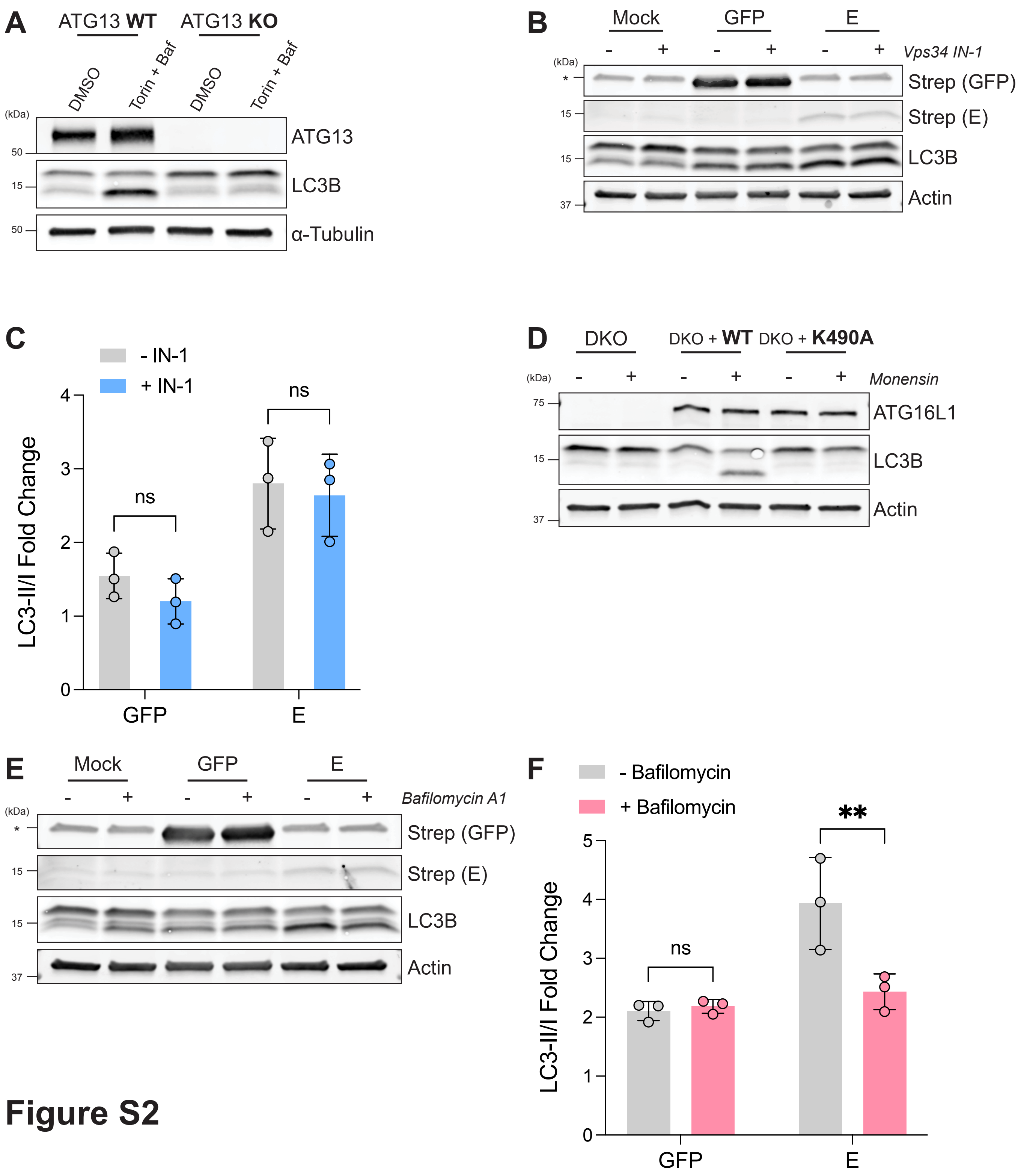

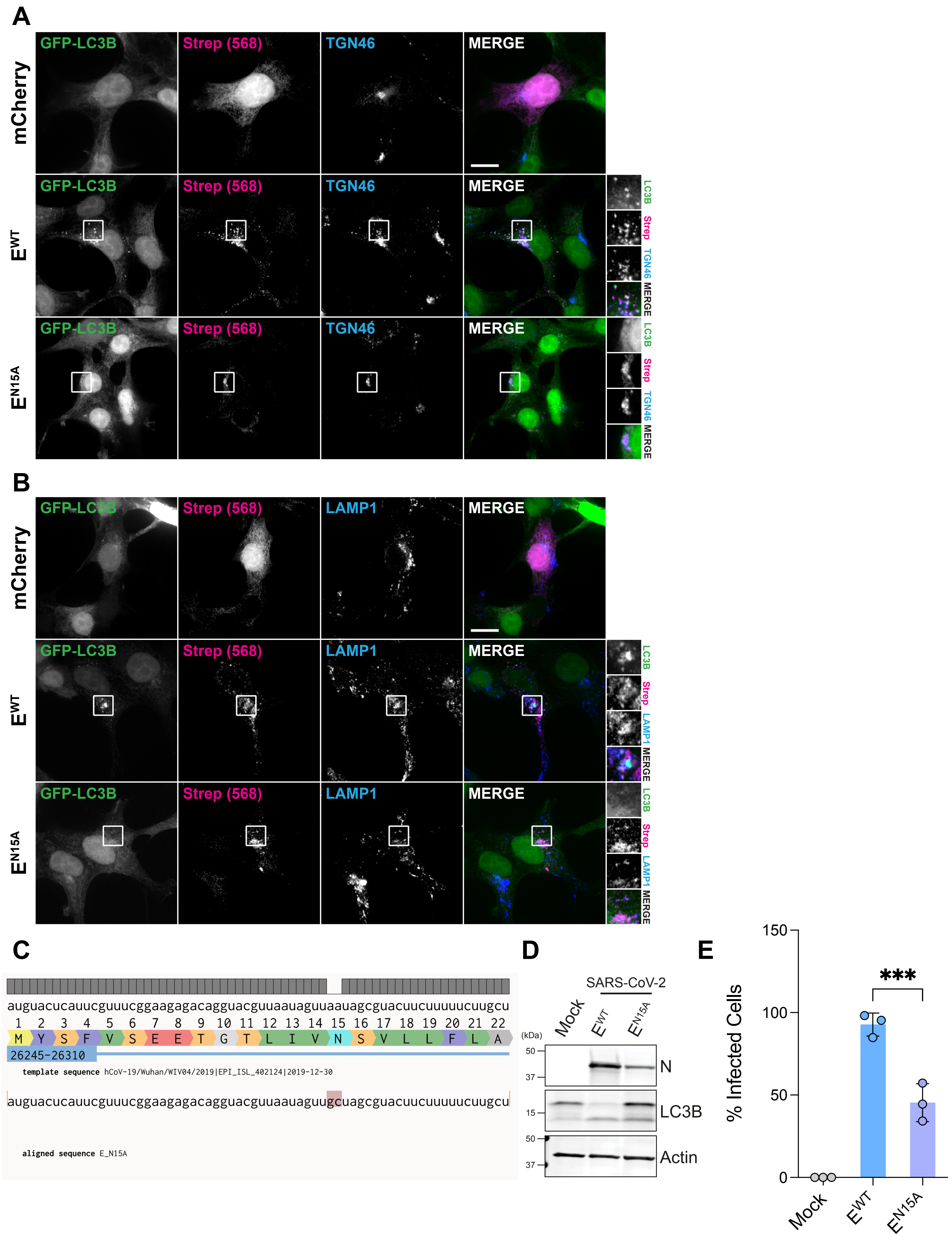
